# *Emx2* Lineage Tracing Reveals Antecedent Patterns of Planar Polarity in the Mouse Inner Ear

**DOI:** 10.1101/2023.10.12.562106

**Authors:** Ellison J. Goodrich, Michael R. Deans

**Affiliations:** Department of Neurobiology, Spencer Fox Eccles School of Medicine at the University of Utah, Salt Lake City, UT 84112, USA; Department of Otolaryngology – Head & Neck Surgery, Spencer Fox Eccles School of Medicine at the University of Utah, Salt Lake City, UT 84132, USA

## Abstract

The planar polarized organization of vestibular hair cells in the utricle and saccule is unique because these inner ear sensory organs contain two groups of hair cells with oppositely oriented stereociliary bundles that meet at a Line of Polarity Reversal (LPR). This organization allows the utricle or the saccule to detect motions directed in opposite directions, and is coordinated with patterns of neural innervation. EMX2 is a transcription factor that is only expressed by hair cells located on one side of the utricle or saccule where it reverses the orientation of their bundles and thereby establishes the position of the LPR. We generated *Emx2*-CreERt2 transgenic mice for genetic lineage tracing and demonstrate robust *Emx2* expression at embryonic day 11.5 (E11.5), before hair cell specification, and when the nascent utricle and saccule have not yet segregated from a common prosensory domain. All hair cells derived from *Emx2*-CreERt2 lineage tracing at E11.5 are restricted to one side of the LPR in the mature utricle or saccule indicating that an antecedent LPR may be established by EMX2 at that stage. Consistent with this, *Emx2*-CreERt2 lineage tracing at E11.5 in *Dreher* mutant mice, where the utricle and saccule fail to segregate, labels a continuous field of cells distributed along one side of a fused utricular-saccular-cochlear organ. Altogether these observations reveal that the origin of the LPR is established in the developing prosensory domain, and that the presence or absence of *Emx2* expression defines progenitor cells with distinct lineages that include hair cells with oppositely oriented stereociliary bundles.

## Introduction

The mature inner ear contains discrete sensory epithelia that detect sound or motion, and mediate auditory and vestibular function (Fig.1A). These include two vestibular organs, the utricle and saccule, which respond to linear acceleration and contain mirrored groups of sensory receptor hair cells capable of detecting accelerations oriented in opposite directions (Fig.1B). Hair cells are specialized mechanoreceptors that extend a bundle of actin-rich stereocilia from their apical surface and transduce microscopic deflections of the stereocilia into neural activity. The stereociliary bundle is structurally and functionally polarized, with the stereocilia arranged in rows of increasing height, and only deflections towards the taller stereocilia open mechanically gated ion channels and depolarize the cell (Fig.1C). A true tubulin-based cilium called the kinocilium is always located adjacent to the tallest row of stereocilia. As a result, the position of the kinocilium and its associated basal body beneath the apical cell surface readily define both the structural and functional polarity of a hair cell.

**Figure 1:**
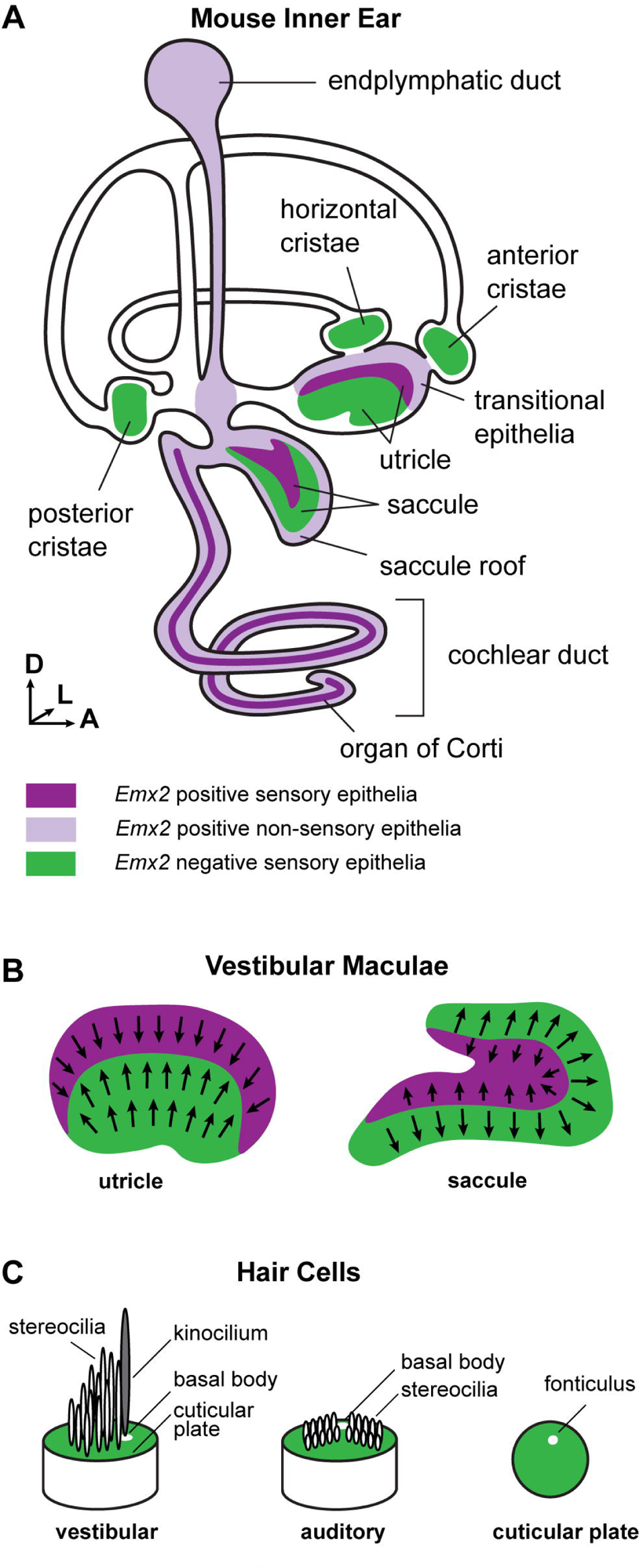
Patterns of *Emx2* expression are correlated with Planar Polarity in the mouse inner ear. (A) Schematic diagram of the membranous labyrinth and sensory epithelia of the mouse inner ear. Magenta shading represents the distribution of *Emx2* expression in sensory (darker shades) and non-sensory tissue (lighter shades). Green illustrates portions of the sensory epithelia that do not express *Emx2*. (B) The LPR is positioned in the vestibular maculae at the boundary between *Emx2*-expressing (magenta) and non-expressing (green) regions. In the utricle hair cells are oriented with the excitatory axis of the bundle pointed towards the LPR while in the saccule the bundles are pointed away. (C) The excitation axis of the hair cell is determined by the organization of the stereocilia in rows of increasing height adjacent to a kinocilium that is retained in vestibular hair cells but lost from auditory hair cells at the onset of hearing. In both cell types a basal body beneath the kinocilium is laterally displaced on the apical cell surface and can be located by the position of the fonticulus.

The stereociliary bundles of utricular and saccular hair cells are embedded in an overlying extracellular matrix called the otoconial membrane, and inertial movements of this membrane during acceleration deflects the stereocilia. The two groups of hair cells in either sensory epithelium have oppositely oriented bundles (Fig.1B) and as a result, movements of the otoconial membrane simultaneously excite one group and inhibit the other group of hair cells. The polarized orientation of the bundle is stable because it is anchored in a cuticular plate composed of filamentous actin extending throughout the apical cell surface with the minor exception of a gap, called the fonticulus, that is occupied by the basal body of the kinocilium (Fig.1C). Head rotation is detected by hair cells in one of three semi-circular canal cristae and sound is detected by hair cells in the organ of Corti of the cochlea (Fig.1A).

This polarized organization of hair cells is called planar polarity because it occurs parallel to the plane of the sensory epithelium, and the orientation of neighboring cells is coordinated by intercellular signaling mediated by the core Planar Cell Polarity (PCP) proteins (Deans 2021). In the utricle and saccule, the two groups of hair cells are adjacent and meet at a cell boundary called the line of polarity reversal (LPR). The position of the LPR is determined by expression of the transcription factor EMX2, and hair cells expressing EMX2 always have stereociliary bundle orientations opposite of those that do not (Fig.1C). Despite this function, EMX2 expression is not restricted to hair cells and occurs throughout sensory and non-sensory structures of the mouse inner ear (Fig.1A). Sensory structures expressing EMX2 include the auditory hair cells and supporting cells of the organ of Corti in the cochlea, and both hair cells and supporting cells along one side of the LPR in the utricle and saccule. Non-sensory structures expressing EMX2 include the remainder of the cochlear duct, the endolymphatic duct, the transitional epithelia of the utricle and epithelial cells comprising the roof of the saccule. EMX2 is not found in the semi-circular canals or their associated cristae (Fig.1A). These patterns of expression have been previously shown by endogenous protein and gene expression and also genetic labeling with *Emx2*-Cre (Kimura, Suda et al. 2005, Holley, Rhodes et al. 2010, Jiang, Kindt et al. 2017, Stoller, Roman et al. 2018, Jia, Ratzan et al. 2023).

EMX2 regulates planar polarity by acting as a master regulator or switch that determines how the stereociliary bundle is oriented relative to an underlying polarity axis established by the core PCP proteins (Deans, Antic et al. 2007, Jiang, Kindt et al. 2017). In *Emx2* mutants the LPR is lost and all of the stereociliary bundles in the utricle or saccule are oriented in the same direction. Conversely, when *Emx2* is overexpressed in hair cells, bundle orientation is completely reversed and the LPR is similarly absent (Jiang, Kindt et al. 2017). EMX2 function is similar in the lateral line neuromasts of zebrafish where *Emx2* is only expressed in half the hair cells and these have the opposite stereociliary bundle orientations as the *Emx2*-negative hair cells (Jiang, Kindt et al. 2017, Lozano-Ortega, Valera et al. 2018, Kozak, Palit et al. 2020). Despite these well characterized effects on hair cell differentiation, *Emx2* expression occurs in the otic vesicle at developmental stages before hair cell specification, and bundle orientation is dependent the polarity effectors GPR156 (Kindt, Akturk et al. 2021) and STK32A (Jia, Ratzan et al. 2023) that are expressed exclusively in hair cells and function downstream of EMX2. The early and broad pattern of *Emx2* expression led us to hypothesize that EMX2 pre-determines the position of the LPR by patterning the prosensory domain shortly after its formation in the otic vesicle, and prior to segregation of the utricle and saccule. This was tested by generating *Emx2*-CreERt2 mice to allow for genetic lineage tracing of cells expressing *Emx2* at early stages of otic development and evaluating their distribution in the mature sensory organs after LPR formation. The potential for the prosensory domain to be patterned by EMX2 was further demonstrated by lineage tracing experiments in the *Dreher* mutant mouse (Nichols, Pauley et al. 2008, Koo, Hill et al. 2009) in which segregation of the utricle and saccule does not occur.

## Results

### Early onset and distribution of *Emx2* expression based upon *Emx2*-Cre genetic labeling

The onset and distribution of *Emx2* gene expression in the developing mouse inner ear can be visualized through genetic labeling by crossing *Emx2* ^Cre/WT^ knock-in mice (*Emx2*-Cre (Kimura, Suda et al. 2005)) with the Cre-dependent reporter ROSA ^tdTom(Ai9)^ (Madisen, Zwingman et al. 2010) to permanently mark *Emx2*-expressing cells with the red fluorescent protein tdTomato. At embryonic day 11.5 (E11.5), *Emx2* ^Cre/WT^; *ROSA* ^tdTom(Ai9)/WT^ embryos show broad expression of tdTomato in the developing otic vesicle when viewed by immunofluorescent labeling of frozen sections (Fig.2). At this stage the otic vesicle has invaginated from the surface ectoderm and neuroblasts destined to form the cochlear-vestibular ganglion are delaminating from the epithelium at the antero-ventral wall of the otic vesicle (Goodrich 2016). TdTomato can be visualize in all sections along the dorsal to ventral axis of the otic vesicle at this stage. In dorsal sections expression is restricted to the endolymphatic duct (Fig.2 A-C). Starting at the point where the endolymphatic duct emerges from the otic vesicle, TdTomato expands posteriorly along the medial wall and broadens in more ventral sections to include cells from both the medial and lateral sides of the otic vesicle (Fig.2 A,D-F). This expression domain extends into the ventral-most region of the otic vesicle that will form the cochlear duct (Fig.2 A,G). At this stage genetic labeling includes all regions of the ventral otic vesicle with the exception of the neurogenic region adjacent to the cochlear-vestibular ganglia.

**Figure 2:**
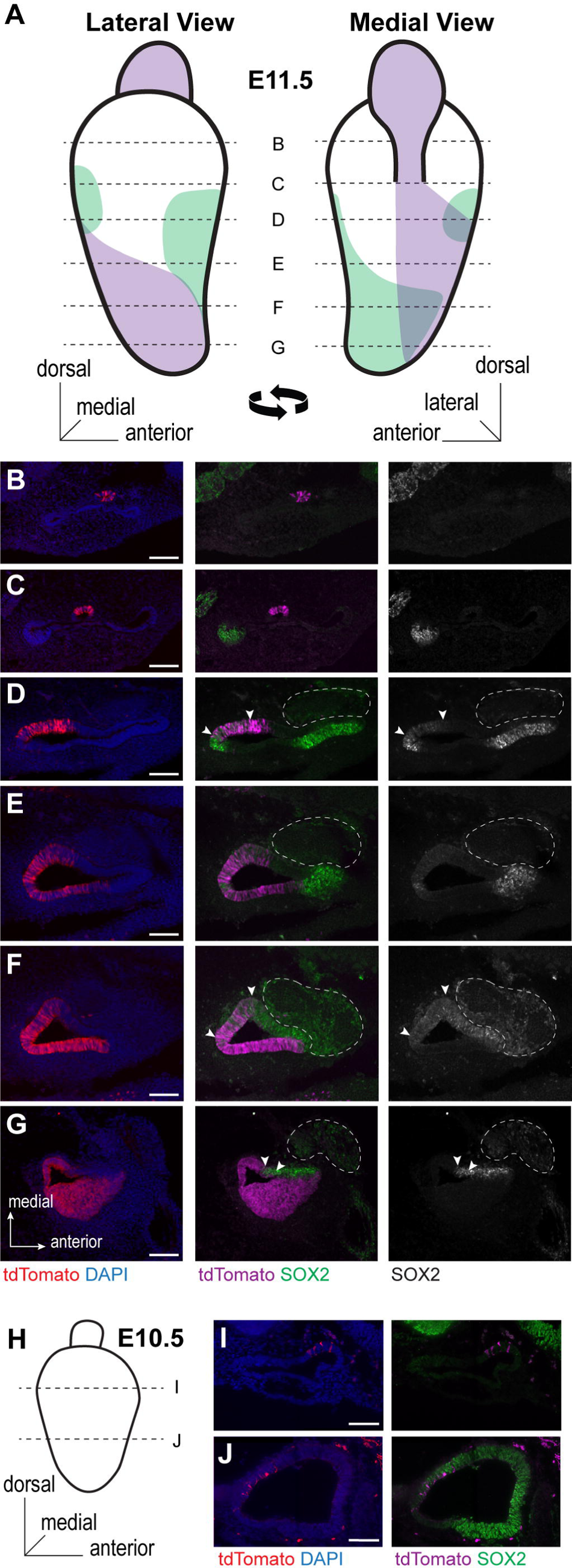
*Emx2*-Cre genetic labeling of the mouse otic vesicle. (A)Schematic diagram of the E11.5 otic vesicle summarizing the distribution of *Emx2*-Cre labeled cells and SOX2 expressing cells based upon immunofluorescent labeling of frozen sections. Dashed lines represent the approximate location of fluorescent images in subsequent panels. (B-G) Representative immunofluorescent images demonstrating *Emx2*-Cre labeled cells and SOX2. Arrowheads mark the boundaries of regions of overlap. (H) Schematic diagram of the E10.5 otic vesicle representing the approximate location of images in subsequent panels. (I-J) *Emx2*-Cre genetic labeling initiates in small groups of cells at E10.5. Scale bars: 100µm

Regions of the otic vesicle with the potential to become sensory epithelia express SOX2, though at E11.5 the lineage of SOX2-expressing cells also includes non-sensory structures (Gu, Brown et al. 2016). Nonetheless SOX2 immunolabeling is a suitable marker for prosensory potential at this stage, and cells genetically labeled by *Emx2*-Cre overlap with SOX2 in two locations. The first is an isolated patch of SOX2-expressing cells on the posterior wall of the otic vesicle that will become the posterior cristae and likely also contributes to the endolymphatic duct (Fig.2D) (Gu, Brown et al. 2016). The second is a region of the medial ventral otic vesicle that extends ventrally through the nascent cochlear duct (Fig.2F,G). This region overlaps with the SOX2-expressing prosensory domain that will give rise to the utricle, saccule and organ of Corti. This is significant because it suggests that cells destined to reside along one side of the LPR in the lateral region of the utricle or inner region of the saccule may already express *Emx2* at this stage. Earlier expression of tdTomato at E10.5 is limited to sparsely distributed cells that do not form a continuous domain (Fig.2H-J). The onset of *Emx2*-Cre recombinase activity at E10.5 has also been observed with a second Cre-dependent reporter line (Ono, Kita et al. 2014) where reporter expression was similarly disperse. Overall, the onset of robust genetic labeling by *Emx2*-Cre at E11.5 coincided with the appearance of the endolymphatic duct, and included both prosensory and non-sensory regions of the otic vesicle.

### Design and Characterization of *Emx2*-CreERt2 mice

To visualize the lineage of cells expressing *Emx2* at E11.5 and determine their distribution relative to the LPR in the mature utricle and saccule, a transgenic mouse expressing tamoxifen-inducible CreERT2 from the endogenous *Emx2* promoter was generated using CRISPR-based gene editing techniques (Fig.3A). The targeting strategy generated a bicistronic *Emx2* mRNA by inserting a P2A sequence between the *Emx2* and the transgenic *CreERt2* coding sequence in order to maintain endogenous EMX2 function. A CRISPR guide RNA was designed to direct a single Cas9 DNA cleavage near the *Emx2* translational stop codon, and donor DNA containing the P2A sequence followed by *CreERt2* replaced the stop codon following DNA repair by homologous recombination. This gene editing approach did not disrupt the *Emx2* coding sequence or gene regulatory sequences, or the 5’ and 3’ untranslated regions (UTRs) of the *Emx2*-*CreERt2* bicistronic mRNA. Founder mice were screened by PCR genotyping using PCR primer pairs flanking the 5’ and 3’ homology regions of the donor DNA (Fig.2B) and a single founder mouse was selected to establish the transgenic line.

**Figure 3:**
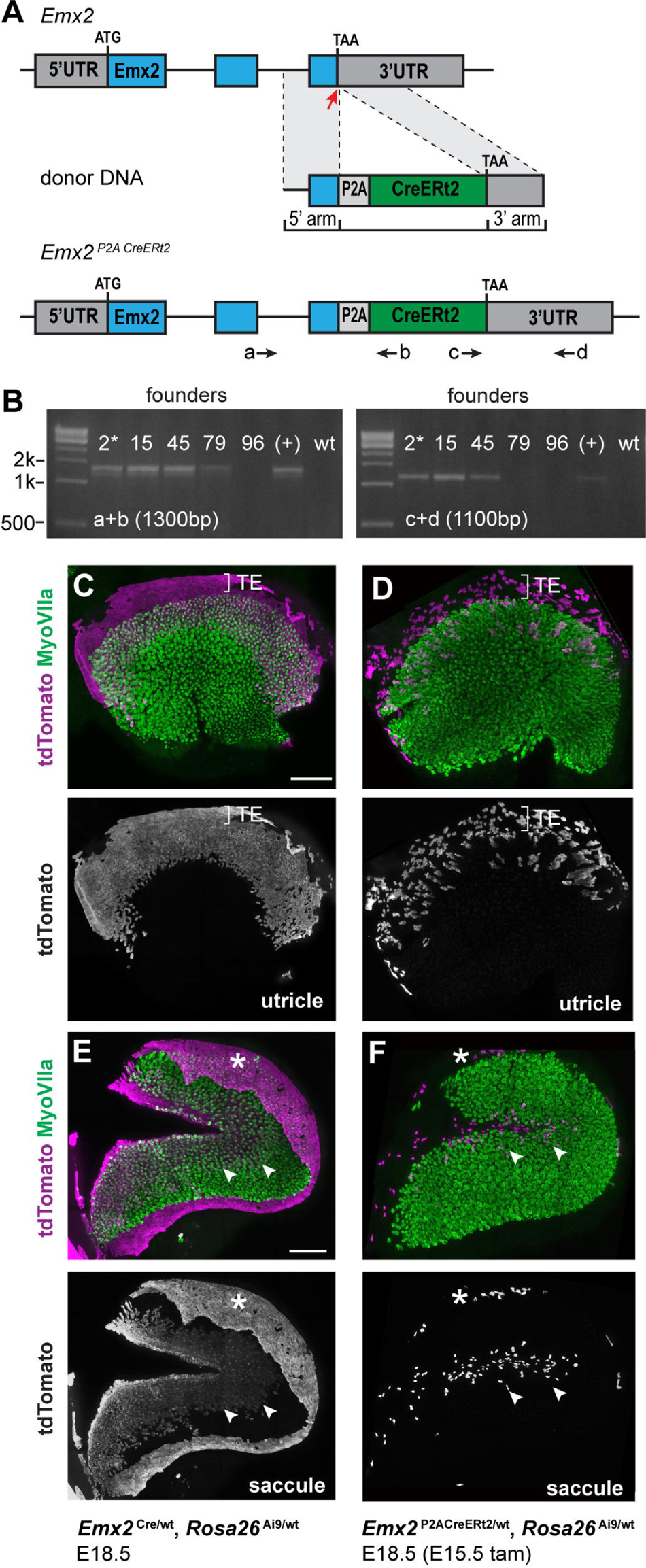
*Emx2*-CreERt2 gene targeting and validation. (A) Crispr-mediated gene targeting strategy used to generate the *Emx2*-CreERt2 line of transgenic mice. Blue marks the *Emx2* coding sequence and the red arrow the position of the Cas9 cut site. (B) PCR-based evaluation of homologous recombination in candidate founders. Positions of PCR primers flanking the 5’ and 3’ homologous arms are illustrated in (A). Mouse #2 was selected to establish the *Emx2*-CreERt2 line. (C) *Emx2*-Cre lineage traced utricle at E18.5 and (D) *Emx2*-CreERt2 lineage traced utricle at E18.5 following activation by a single dose of tamoxifen delivered at E15.5. Bracket denotes boundaries of the Transitional Epithelia (TE). (E) *Emx2*-Cre lineage traced saccule at E18.5 and (F) *Emx2*-CreErt2 lineage traced saccule at E18.5 following activation by a single dose of tamoxifen delivered at E15.5. Asterisks mark labeled cells in the saccular roof where this tissue was not removed during dissection. Arrowheads mark a boundary of labeled cells within the saccular sensory epithelium. Scale bars: 100µm

Lineage tracing using the *Emx2* ^Cre^ knock-in line provides a cumulative readout of *Emx2* transcription during the course of inner ear development when crossed with the tdTomato reporter line Ai9 and evaluated at perinatal stages (Kimura, Suda et al. 2005, Madisen, Zwingman et al. 2010, Jiang, Kindt et al. 2017, Stoller, Roman et al. 2018). Within the utricle and the saccule this pattern includes regions of the sensory epithelia located along one side of the LPR (Fig.3C,E). Also labeled are non-sensory epithelia located adjacent to the vestibular hair cells. These are the transitional epithelia of the utricle (Fig.3C) and epithelial cells comprising the roof of the saccule (Fig.3E). A similar pattern of genetic labeling is seen in *Emx2* ^P2ACreERt2/wt^; *ROSA* ^Ai9/wt^ mice following tamoxifen-induced activation of CreERt2 at E15.5 (Fig.3D,F). In contrast to Cre recombinase, genetic labeling by CreERt2 only marks cells expressing the transgene for a short period of time following tamoxifen induction, estimated to be 24 hours, and the number of cells labeled depends upon tamoxifen dose and levels of transgene expression. The fewer number of cells labeled by *Emx2*-CreERt2 compared to *Emx2*-Cre likely reflects the levels of CreERt2 expression from the *Emx2* locus since the number of labeled cells is increased in *Emx2* ^P2ACreERt2/P2ACreERt2^ embryos treated with the same amount of Tamoxifen (Fig.5L,M). Larger doses of Tamoxifen proved detrimental to the developing litters which were frequently miscarried despite the inclusion of Progesterone to mitigate this negative effect of Tamoxifen treatment.

A consequence of the knock-in strategy employed to generate the *Emx2*-Cre line is that *Emx2* ^Cre/Cre^ mice have mutant phenotypes similar to those reported for other *Emx2* mutants (Kimura, Suda et al. 2005, Holley, Rhodes et al. 2010, Jia, Ratzan et al. 2023). In these mice the LPR does not form because vestibular hair cells in the lateral region of the utricle are misoriented and share the same bundle orientations as hair cells in the medial region. In contrast, LPR formation occurs normally in the *Emx2* ^P2ACreERt2/P2ACreERt2^ utricle (Fig.4A).

**Figure 4:**
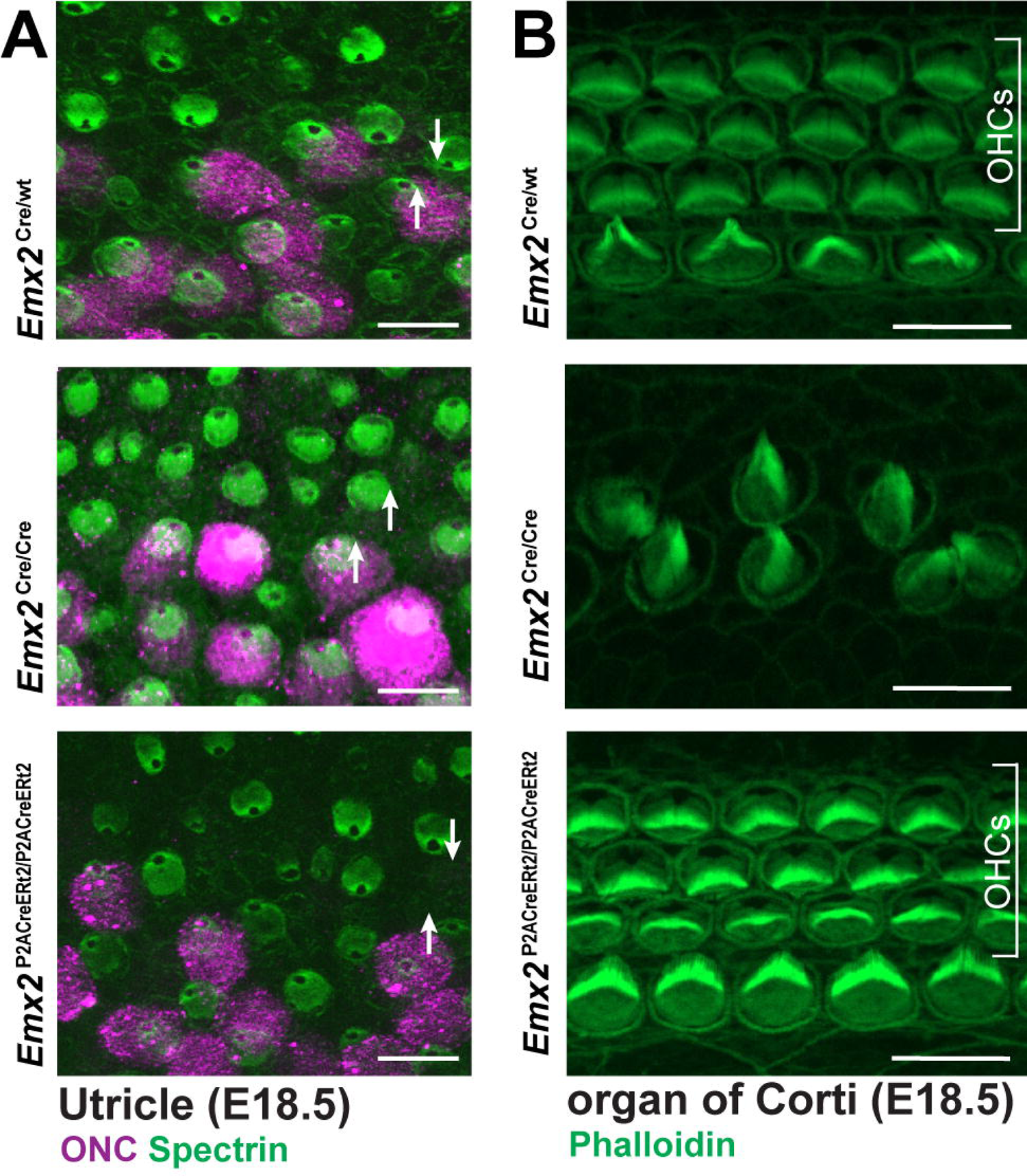
The *Emx2*-CreERt2 transgene does not disrupt EMX2 function. (A) The organization of vestibular hair cells and patterning about the LPR in the utricle of heterozygous control, or *Emx2*-Cre and *Emx2*-CreERt2 transgenic mice when the transgenes are homozygosed. βII-Spectrin labels the hair cells and reveals bundle orientation based upon the position of the fonticulus. Oncomodulin (ONC) labels hair cells in the striolar region which is located along one side of the LPR. Arrows indicate bundle orientation for representative pairs of hair cells. (B) The organization of auditory hair cell stereociliary bundles in the organ of Corti of heterozygous control, or *Emx2*-Cre and *Emx2*-CreERt2 transgenic mice when the transgenes are homozygosed. OHC (Outer Hair Cells). Scale bars: 10µm

*Emx2* is also required for outer hair cell (OHC) development in the mouse cochlea (Holley, Rhodes et al. 2010) and OHCs are similarly missing, and the patterning and planar polarity of inner hair cells (IHC) is disrupted, in *Emx2* ^Cre/Cre^ mice (Fig.4B). In contrast, all three rows of OHCs and the IHCs are intact and organized with normal planar polarity in *Emx2* ^P2ACreERt2/P2ACreERt2^ cochlea (Fig.4). Together these phenotypes demonstrate that *Emx2* expression is not disrupted in in the *Emx2*-CreERt2 line where levels of EMX2 are sufficient to support normal inner ear development.

### *Emx2* Lineage Tracing Throughout Inner Ear Development

The sensory epithelia of the inner ear are derived from a common prosensory domain that is specified within the otic vesicle shortly after invagination and fusion at E10 (Mann, Galvez et al. 2017, Zak and Daudet 2021). At this stage the otic vesicle is also patterned about its major axes by morphogen gradients emanating from peri-otic tissues (Bok, Chang et al. 2007, Bok, Dolson et al. 2007, Whitfield and Hammond 2007). Based upon these patterns, the prosensory domain is segregating into distinct auditory and vestibular domains, and subsequently into separate sensory organ precursors. The first to appear is the posterior cristae followed shortly after by the remaining 4 vestibular sensory epithelia and the organ of Corti (Morsli, Choo et al. 1998).

As a result, the utricle, saccule and anterior and horizontal canals share a common origin while the posterior cristae is independently specified. The early expression of *Emx2*-Cre (Fig.2) suggests that the prosensory region might also be pre-patterned into sub-domains that determine the position of the LPR along their boundaries at later stages. This hypothesis was tested by lineage tracing cells within the developing ear using the *Emx2*-CreERT2 line.

To facilitate genetic lineage tracing, *Emx2* ^P2ACreERt2/wt^ males were crossed with Rosa^Ai9/Ai9^ females and a single dose of tamoxifen supplemented with progesterone was delivered by gavage to timed pregnant mice at E10.5, E11.5, E12.5 or E13.5 and the extent of Cre-mediated lineage tracing evaluated at P0 (Fig.5B). For these experiments it is presumed that CreERt2 is activated by tamoxifen for a 24-hour window. Consistent with frozen sections of *Emx2* ^Cre/wt^; Rosa ^Ai9/wt^ otic vesicles at E10.5, tamoxifen induction of *Emx2*-CreERt2 at E10.5 did not label cells within the utricle and saccule when evaluated at P0 (Fig.5C,D). However, a single dose of tamoxifen delivered at E11.5 was sufficient to label cells within the sensory epithelia of the utricle and the adjacent transitional epithelia (Fig.5C). Sparse labeling of cells within the inner region of the saccule was also seen at this stage (Fig.5D). Labeling of both the utricle and saccule was more robust when tamoxifen was delivered at E12.5 and was consistently seen following induction at later stages (Fig.3D,F and Fig.5C,D). For more robust genetic labeling, the *Emx2*-CreERt2 allele was homozygosed and the lineage of *Emx2* expressing cells was evaluated at P0 following tamoxifen induction at E10.5 or E11.5 in *Emx2* ^P2ACreERt2/P2ACreERt2^; *Rosa* ^Ai9/wt^ embryos. Patterns of labeled cells were consistent with the onset of *Emx2* expression and occurred in a subset of cells labeled at E10.5 and expression throughout sensory cells located along one side of the LPR in both the utricle and saccule when cells were labeled at E11.5 (Fig.5E,F). This labeling protocol also marked non-sensory cells in the transitional epithelia and saccule roof in a pattern similar to the cumulative labeling seen with *Emx2*-Cre.

**Figure 5:**
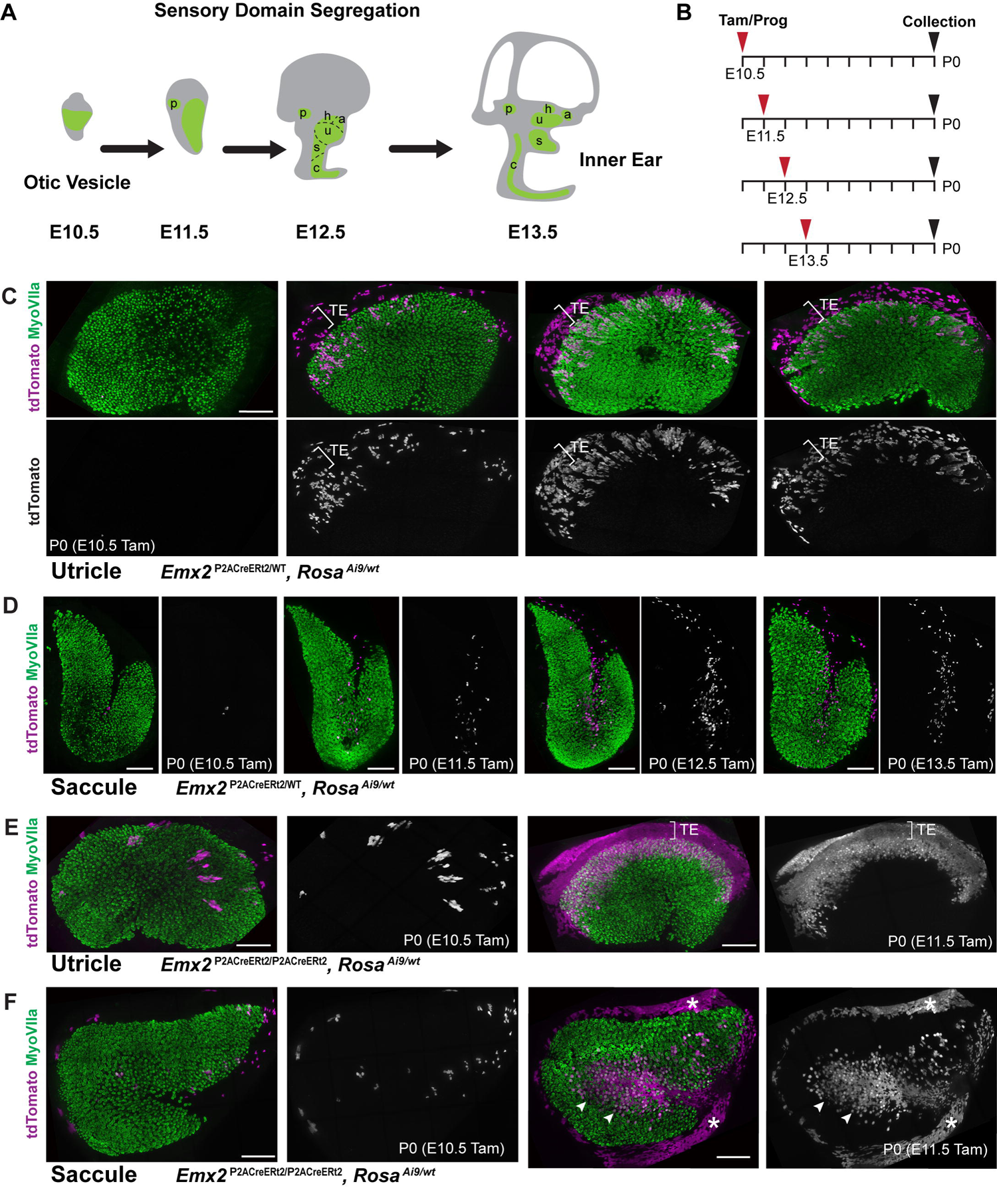
The *Emx2*-CreERt2 lineage is established prior to segregation of the vestibular maculae. (A) Schematic illustrating sensory domain segregation in the mouse inner ear. The sensory epithelia are derived from a common prosensory domain (green) that is segregated over time to form the posterior (P), horizontal (h) and anterior (a) semi-circular canal cristae, the utricle (u) and saccule (s), and the organ of Corti (c). (B) Experimental timeline for *Emx2*-CreERt2 lineage tracing experiments illustrated in subsequent panels. (C-D) tdTomato expression in P0 *Emx2* ^P2ACreERt2/wt^; Rosa ^Ai9/wt^ utricles and saccules following a single dose of tamoxifen delivered between E10.5 to E13.5 as indicated. Hair cell labeling with MyoVIIa antibodies defines the boundaries of the sensory epithelia and bracket indicates the approximate width of the transitional epithelia. (E-F) tdTomato expression in P0 Emx2 ^P2ACreERt2/P2ACreERt2^; Rosa ^Ai9/wt^ utricles and saccules following a single dose of tamoxifen delivered at E10.5 or E11.5. Arrowheads indicate the boundary of the LPR in the saccule and Asterisks mark labeling of the saccular roof. Scale bars: 100µm

The distribution of sensory and non-sensory cells labeled by *Emx2*-CreERt2 was further evaluated by immunofluorescent labeling of frozen sections of P0 ears following tamoxifen induction at E11.5 (Fig.6A). Consistent with wholemount imaging, tdTomato was seen in hair cells and supporting cells of the lateral region of the utricle and transitional epithelia cells (Fig.6B), hair cells and supporting cells within the saccule and non-sensory cells throughout the roof of the saccule (Fig.6C). tdTomato was also present throughout the cochlear duct including the greater epithelial ridge (GER), hair cells within the organ of Corti, and cells throughout the lateral wall (Fig.6D) in addition to the endolymphatic duct (Fig.6E). Outside of the inner ear, *Emx2*-CreERt2 lineage tracing also revealed *Emx2* expression in cells contributing to the ossicular chain (Fig.6A) which is consistent with the role of this transcription factor in their development, and the hearing loss reported in the *Emx2* mutant mouse *Pardon* resulting from ossicular malformations (Rhodes, Parkinson et al. 2003).

**Figure 6:**
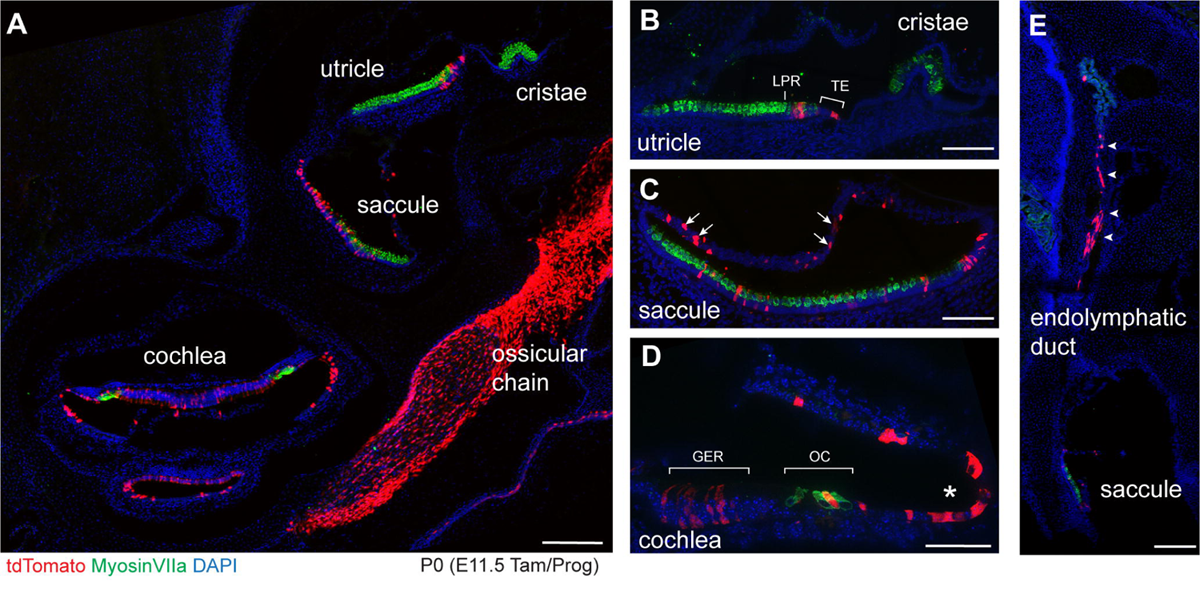
The *Emx2*-CreERt2 lineage following tamoxifen induction at E11.5. (A) Cross section through a *Emx2* ^P2ACreERt2/wt^; Rosa ^Ai9/wt^ inner ear at P0 that had received a single dose of tamoxifen at E11.5. Reporter expression is evident in the utricle, saccule and cochlea, in addition to robust labeling of the ossicular chain. No labeling is seen in the cristae. (B) Cross-section of the utricle showing tdTomato labeled cells along one side of the presumptive LPR and the adjacent transitional epithelia (TE). (C) Cross-section of the saccule showing tdTomato labeled cells within the sensory epithelia as well as non-sensory cells located in the epithelia that comprises the roof of the saccule (arrows). (D) A cross-section of one turn of the cochlea showing labeled cells within the greater epithelial ridge (GER), organ of Corti, and non-sensory cells within the lateral wall of the cochlear duct (asterisk). (E) Labeled cells are also detected within the endolymphatic duct (arrowheads). Scale bars: 200µm (A,E), 100µm (B,C), 50µm (D)

Although the efficiency of Cre-mediated recombination in *Emx2*-CreERt2 mice was low, this level of genetic labeling does enable easy quantification of the frequency distribution of cell types that were labeled at each developmental stage (Fig.7A,B). Within the utricle, tamoxifen induction at E11.5 resulted in a comparable number of hair cell, supporting cell and transitional epithelial cells labeled. However, the frequency of hair cell labeling peaked at E12.5, and at subsequent stages tamoxifen induction was more likely to result in the labeling of supporting cells and transitional epithelia cells. This difference is unlikely to be the result of cell proliferation since labeling at E11.5 that would preferentially mark supporting cells and transitional epithelia which are continually dividing while hair cells are post-mitotic. Instead these labeling patterns are more likely to reflect greater expression of *Emx2* in these different cell types at later developmental stages. Finally, at E15.5 lineage tracing of transitional epithelial cells surpasses that of hair cells and supporting cells within the sensory epithelia. Although it lacks transitional epithelia, a similar transition occurs in the developing saccule with hair cell labeling greatest upon Cre-activation at E11.5 and supporting cells labeled to a greater extent at later ages (Fig.7C,D).

**Figure 7:**
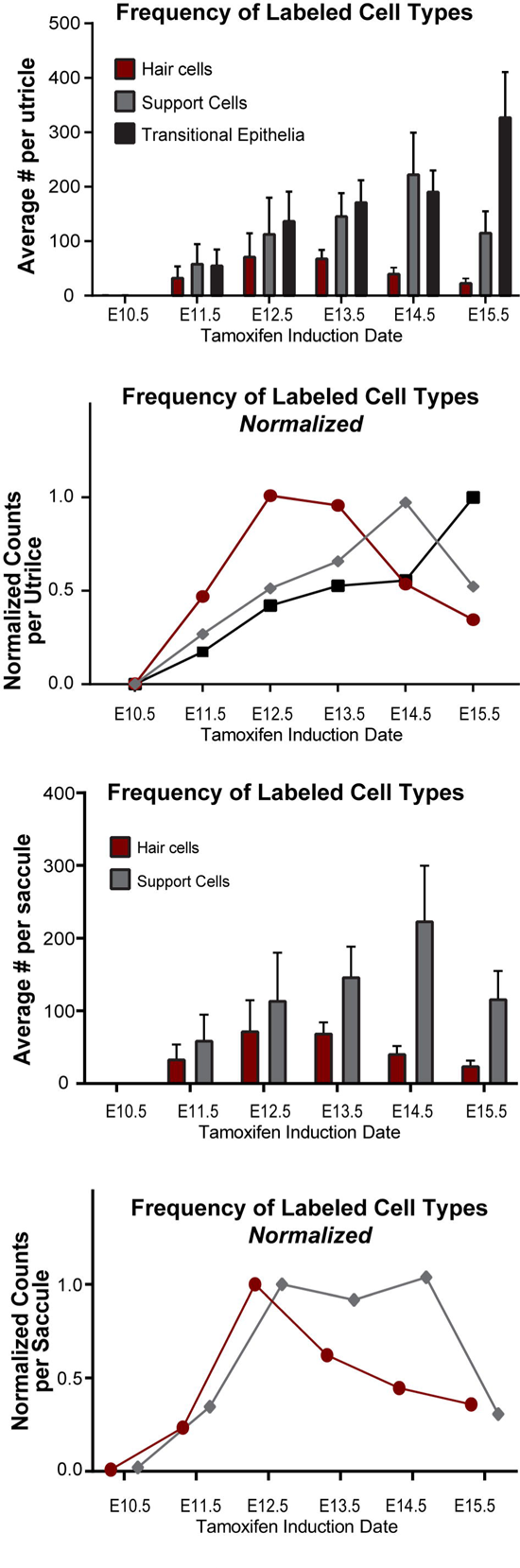
The frequency of labeled cell types is dynamic and changes across development stages. (A) The average number of cells of each type labeled in the *Emx2* ^P2ACreERt2/wt^ utricle at P0 following tamoxifen induction at a single developmental stage, and (B) the relative number of cells of each type normalized to the developmental stage of maximum labeling. n=6 11.5 utricles, n=9 E12.5, n=5 E13.5, n=5 E14.5, n=8 E15.5 (C) The average number of hair cells and supporting cells labeled in the *Emx2* ^P2ACreERt2/wt^ saccule at P0 following tamoxifen induction at a single developmental stage, and (D) the relative number of cells of each type normalized to the developmental stage of maximum labeling. n=3 11.5 saccules, n=9 E12.5, n=4 E13.5, n=4 E14.5, n=6 E15.5

### *Emx2* Expression in the Otic Prosensory Domain Precedes LPR Formation in the Vestibular Maculae

One striking line of evidence that the utricle, saccule and organ of Corti are derived from a common prosensory domain is the mutant phenotype seen in mouse mutants impacting the transcription factor LMX1A including the *Dreher*, *Mutanlallemand*, and *Belly Spot and Deafness* lines (Nichols, Pauley et al. 2008, Koo, Hill et al. 2009, Steffes, Lorente-Canovas et al. 2012). In *Dreher* mutants, the non-sensory cells forming structures that separate the individual sensory epithelia fail to be specified or to proliferate resulting in a continuous utriculo-saccular-cochlear organ in which presumptive macular and cochlear organs are contiguous. Development of the semi-circular canal cristae are also disrupted in the *Dreher* mutants but instead of being fused with the maculae, they form extensive large and branched sensory epithelia. These aberrant cristae are easily identified based upon Prox1 expression and their organization is highly variable between individual mice. In contrast features of the fused utriculo-saccular-cochelar organ are more consistent between individual *Dreher* mutants (Fig.8A).

**Figure 8:**
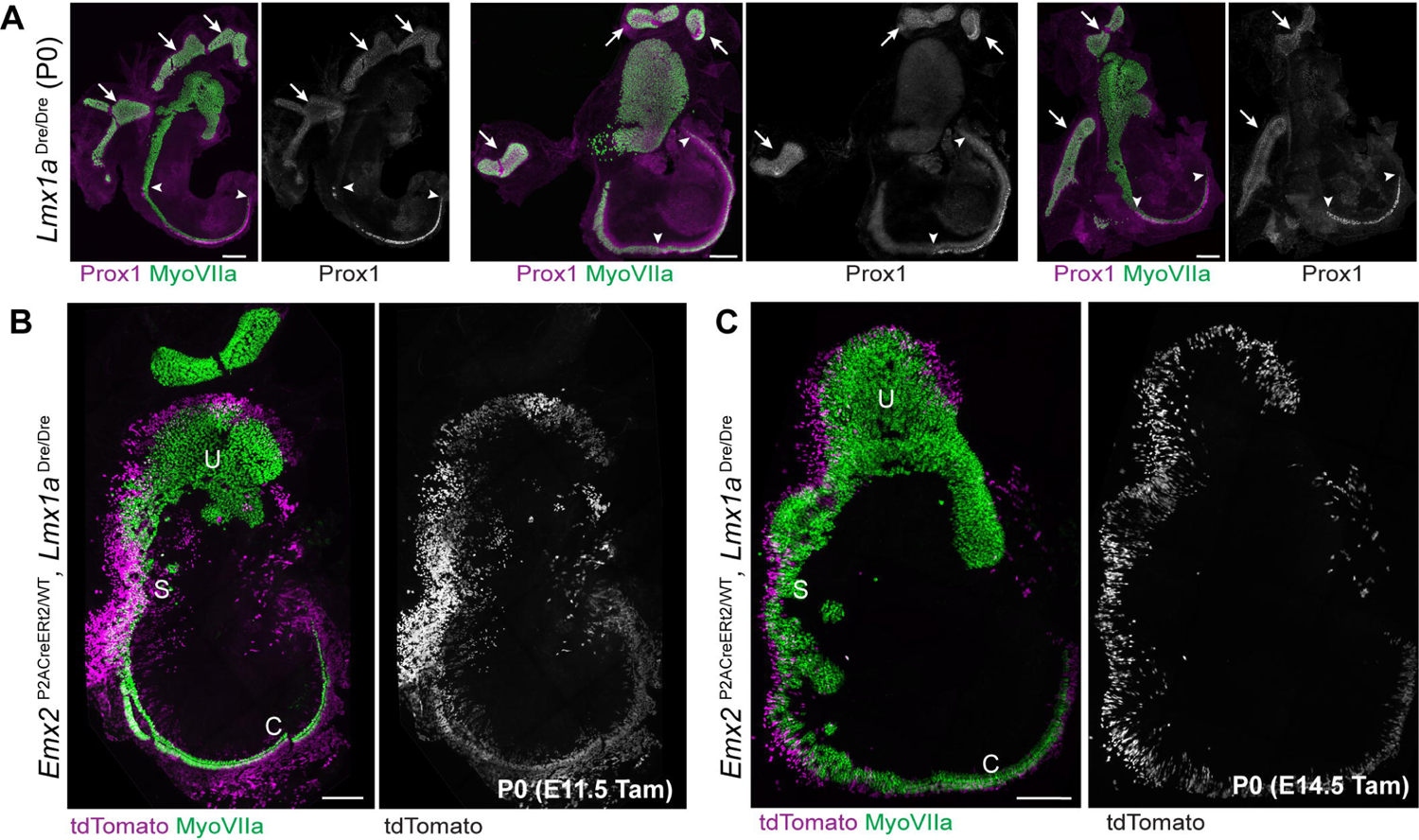
*Emx2*-CreERt2 lineage tracing in *Dreher* mutants. (A) Three examples of the *Dreher* mutant phenotype in which the utricle (U), saccule (S) and cochlea (C) fail to segregate and form a continuous field of hair cells. The transcription factor Prox1 marks auditory hair cells and distinguishes them from hair cells in the presumptive saccule and utricle. Auditory hair cells are bounded by arrowheads. Prox1 also marks vestibular hair cells in the cristae (arrows) whose morphogenesis is highly variable between individual mutants. (B) *Emx2*-CreERt2 lineage tracing in a *Dreher* mutant at P0 following tamoxifen induction at E11.5 labels a field of cells spanning the presumptive utricle (U) and saccule (S). (C) *Emx2*-CreERt2 lineage tracing in a *Dreher* mutant at P0 following tamoxifen induction at E14.5 reveals a similar field of labeled cells spanning the fused utricle and saccule. Scale bars: 200µm

The early expression of *Emx2* suggests that EMX2 could pre-pattern the prosensory domain prior to sensory patch segregation. Since *Emx2*-CreERt2 induction at E11.5 marks cells in the mature utricle and saccule, *Emx2*-CreERt2 lineage tracing of *Dreher* mutants at this stage should reveal the antecedent pattern of *Emx2* expression and a continuous field of tdTomato-labeled cells is expected in the Dreher mutants. This possibility was tested by intercrossing the *Emx2*-CreERt2 and *Dreher* lines and administering tamoxifen at E11.5.

Consistent with this hypothesis, tamoxifen induction at this stage results in a single and continuous field of tdTomato-labeling in Dreher mutants when evaluated at P0 (Fig.8B). This field extends along the boundary of the presumptive utricle and saccule and into the mutant cochlea. Induction at E14.5, a stage in which the LPR would have already formed in wild type mice, results in a similar distribution of tdTomato labeled cells in the *Dreher* mutant ear (Fig.8C). Thus, the lineage of cells expressing EMX2 is established at the earliest stages of prosensory domain development and therefore the origin of the LPR is determined prior to the formation of independent and structurally distinct vestibular maculae.

## Discussion

Using Cre-ERt2 mediated genetic lineage tracing we demonstrate that the planar polarity transcription factor EMX2 is expressed in the prosensory domain of the developing inner ear prior to hair cell specification and before segregation of the utricle and saccule into discrete sensory organs. In addition, we show that the lineage of *Emx2*-expressing cells from this stage is restricted to one side of the LPR in the mature sensory organ. Together these data support the hypothesis that planar polarity signals are established at early stages of inner ear development where they are pre-positioned to guide subsequent steps of hair cell differentiation including stereociliary bundle polarization. These observations were made using a new mouse line in which CreERt2 is expressed and regulated by endogenous *Emx2* gene regulatory elements. Based upon patterns of Cre-mediated genetic labeling, the expression of *Emx2*-CreERt2 matches *Emx2*-Cre and patterns of *Emx2* expression determined by immunolabeling and ISH (Holley, Rhodes et al. 2010, Jiang, Kindt et al. 2017, Stoller, Roman et al. 2018, Jia, Ratzan et al. 2023). However, unlike *Emx2*-Cre, the *Emx2*-CreERt2 line does not disrupt EMX2 function as demonstrated by the lack of mutant phenotypes when the transgene is homozygosed. An unfortunate deficit of *Emx2*-CreERt2 is the limited efficiency of Cre-mediated recombination when tamoxifen is delivered by gavage since that hinders its usefulness for generating conditional knockout mice. We currently have not determined whether *Emx2*-CreERt2 activation is more responsive to intraperitoneal (IP) delivery of 4-hydroxy Tamoxifen, or if CreERt2 induction is more efficient in other tissues where expression from the *Emx2* promoter might be stronger.

### Patterning the developing prosensory domain

The utricle and saccule are derived from adjacent regions of the prosensory domain that have not segregated at E11.5 when *Emx2*-CreERt2 lineage tracing marks cells in both sensory epithelia. Moreover, the early pattern of *Emx2* expression overlaps with the prosensory domain marker SOX2. These results indicate that the prosensory domain is being functionally patterned in the developing otocyst and that patterning does more than distinguish auditory from vestibular and maculae from cristae, but also pre-determines functionality of the future sensory epithelia. Fritzsch and colleagues had suggested that this may be the case when they proposed that the cochlea shared functional lineage with portions of the utricle and saccule based upon evolutionary observations (Fritzsch, Beisel et al. 2002). We propose that these lineages are distinguished by expression of *Emx2* and that the antecedent patterning of the prosensory domain is reflected in the *Emx2*-CreERt2 lineage of *Dreher* mutants (Fig.8). In this scenario, the *Emx2*-expressing domain that overlaps with SOX2 in the ventro-medial otic vesicle (Fig.2) contributes to the utricle and saccule and the boundary of *Emx2* expression is the nascent LPR.

A limitation to the current study is that it consists of snapshots of single developmental stages and cannot track the dynamic morphogenesis linking *Emx2*-Cre and SOX2 overlap at E11.5 with EMX2 expression in the mature utricle and saccule. While imaging these processes in the living embryo is not possible, a detailed timecourse may be practical to construct using light sheet microscopy and 3D renderings of the inner ear if they are collected across a range of developmental stages. Not only would this morphological progression reveal how the saccule is derived from the otic vesicle, it has the potential to explain how a boundary of *Emx2* expression in the E11.5 otic vesicle could form LPRs in the utricle and saccule with opposite organizations of stereociliary bundles.

An outstanding question is what determines the boundaries of EMX2 expression at this stage since this pattern is correlated with the position of the LPR in the mature maculae. The only region of the ventral otic vesicle that is not genetically labeled by *Emx2*-Cre at E11.5 is the antero-medial segment adjacent to the cochleovestibular ganglia (Fig.2). This corresponds to the neurogenic domain from which neuroblasts actively delaminate to form the neighboring ganglia. One possibility is that the neurogenic progression occurring in this domain is incompatible with EMX2 and the allele is actively repressed. It is also possible that a morphogenic signal released from the newly formed ganglia contributes to otic vesicle patterning by repressing *Emx2*-expression in this adjacent region. In either event, this repression appears to be incomplete since a few spiral ganglion neurons are labeled by *Emx2*-Cre (Ghimire, Ratzan et al. 2018) which could be neuroblasts that transiently expressed *Emx2* before delaminating. Alternatively, the boundaries of *Emx2* expression are determined by a combination of morphogenic gradients that are known to establish the primary axes of the otic vesicle at this stage (Bok, Chang et al. 2007). Since boundaries of EMX2 expression are associated with the position of the LPR in mature utricle and saccule, the molecular mechanism(s) that prevent *Emx2*-expression in this region may have a significant impact on vestibular planar polarity in the mature maculae and therefore would be important to identify.

It has also been shown that the expression of the Core PCP proteins that coordinate the orientation of stereociliary bundles between neighboring hair cells is similarly initiated at these early stages of otic development and is maintained through the course of inner ear morphogenesis (Yang, Qian et al. 2017). Based upon similar ideas we had proposed a model in which a restricted pattern of *Emx2* expression and a uniform field of PCP signaling could organize the utricle and the saccule prior to their segregation and account for the complementary patterning about their LPRs. In this theoretical model an annular distribution of EMX2, if divided properly during segregation, would account for the opposite organization of hair cells at the LPR in both maculae (Deans 2013). Of course, the distribution of *Emx2* lineage tracing we see in *Dreher* mutants is not annular, and the final organization of hair cells in the utricle and saccule is dependent upon morphogenesis that does not occur in *Dreher* mutants. Nonetheless, if these two essential organizing principles were established early enough during development, this would only need to occur once and not independently within each sensory organ. Establishing planar polarity patterns early may also account for how different species have evolved alternative sensory structures that share conserved planar polarity features including the coordinated orientation of stereociliary bundles and LPRs.

### Does EMX2 strictly regulate regional patterning?

Within the inner ear and lateral line EMX2 is been well described as a genetic switch that regulates stereociliary bundle polarity (Jiang, Kindt et al. 2017, Ji, Warrier et al. 2018, Kozak, Palit et al. 2020). Thus, when Emx2 is selectively deleted from hair cells their bundles are reoriented (Ji, Tona et al. 2022) or when Emx2 is selectively over-expressed in hair cells the LPR is lost because all bundles have the same orientation (Jiang, Kindt et al. 2017). We have shown that EMX2 functions by negatively regulating the kinase gene *Stk32a*, thereby restricting STK32A expression to hair cells in the medial region of the utricle and outer region of the saccule. Through its kinase activity, STK32A negatively regulates the localization of the planar polarity receptor GPR156 at apical cell boundaries. Due to this double-negative regulatory circuit, GPR156 can only reorient stereociliary bundles relative the underlying PCP axis in cells expressing EMX2 (Kindt, Akturk et al. 2021, Jia, Ratzan et al. 2023).

Despite this recent progress, EMX2 may be better known as a transcription factor that regulates regional identity in the developing neocortex (Bishop, Goudreau et al. 2000, Mallamaci, Muzio et al. 2000). In the absence of *Emx2*, the size of primary cortical regions changes with reductions in rostral areas and corresponding increases in caudal areas whereas upon overexpression of *Emx2* these trends are reversed. This process requires cross-talk between mutually antagonistic transcription factors and is a process known as arealization (Bishop, Goudreau et al. 2000, Hamasaki, Leingartner et al. 2004). Within the inner ear, other features of the *Emx2* mutant phenotype and patterns of *Emx2* expression suggest that it may regulate regional patterning in this context as well. First, EMX2 is not restricted to hair cells like other transcription factors required for hair cell development (Bermingham, Hassan et al. 1999, Keithley, Erkman et al. 1999, Wallis, Hamblen et al. 2003), and can be detected in non-sensory structures including the transitional epithelia of the utricle, roof of the saccule and the endolymphatic duct. Second, in *Emx2* mutants the most prominent cochlear phenotype is a loss of OHCs and severe IHC disorganization rather than a reversal of stereociliary bundle orientation (Holley, Rhodes et al. 2010) (also Fig.4B). Finally, as we have demonstrated, *Emx2* transcription initiates at a time when the otocyst is actively being patterned along its primary axes.

One possibility is that the primary function of EMX2 is to establish the regional identity of sensory epithelia, dividing the utricle and saccule into distinct functional domains with bundle orientation being the most salient indication that patterning has occurred. In this case we predict that EMX2 would sit atop a gene regulatory network that patterns the utricle and saccule and thus there are likely to be additional factors acting between EMX2 and STK32A. Alternatively, it is possible that EMX2 has multiple functions and that after otocyst patterning, it acts in hair cells to regulate factors such as STK32A required to orient the stereociliary bundle. This would be consistent with hair cell overexpression studies where the EMX2 effect is direct and cell specific (Jiang, Kindt et al. 2017). Distinguishing between these alternatives would reveal whether or not there are additional planar polarity factors waiting to be discovered or if all of the factors regulating vestibular hair cell polarization have been identified.

## Methods

### Mouse lines and husbandry

All mice were housed at the University of Utah under Institutional Animal Care and Use Committee (IACUC) approved guidelines, and individual lines were maintained by backcross to B6129SF1/J hybrid females and genotyped using allele-specific PCRs (available upon request). For timed breeding and tissue staging, noon on the day in which a vaginal plug was seen was considered embryonic day 0.5 (E0.5), and the day litters were born was considered postnatal day 0 (P0). Mice from both sexes were used for experimentation. *Emx2* ^Cre/WT^; *Rosa* ^tdTomAi9/WT^ or *Emx2* ^P2ACreERt2/WT^; *Rosa* ^tdTomAi9/WT^ embryos were generated by crossing *Emx2* ^Cre/WT^, *Emx2* ^P2ACreERt2/WT^ or *Emx2* ^P2ACreERt2/P2ACreERt2^ male mice with *Rosa* ^tdTomAi9/tdTomAi9^ females and staged and collected as described. *Emx2* ^P2ACreERt2/P2ACreERt2^; *Rosa* ^tdTomAi9/WT^ embryos were generated by crossing *Emx2* ^P2ACreERt2/P2ACreERt2^ male and *Emx2* ^P2ACreERt2/WT^; *Rosa* ^tdTomAi9/WT^ female mice. *Emx2* ^Cre/Cre^ embryos and *Emx2* ^P2ACreERt2/P2ACreERt2^ pups were generated by intercrossing male and female heterozygotes for each allele. *Lmx1a Dreher* mutants were generated by intercrossing *Lmx1a* ^Dre/WT^ males and females while for *Emx2*-CreERt2 lineage tracing *Emx2* ^P2ACreERt2/P2ACreERt2^; *Lmx1a* ^Dre/WT^ males were crossed with *Lmx1a* ^Dre/WT^; *Rosa* ^tdTomAi9/tdTomAi9^ females. *Emx2*-Cre mice (Kimura, Suda et al. 2005) were provided by S. Aizawa (Riken Institute). B6129SF1/J hybrid (strain #101043), *Lmx1a/Dreher* (strain #000636) and *Rosa* Ai9tdTOM (strain #007909) (Madisen, Zwingman et al. 2010) mice were purchased from The Jackson Laboratory.

### Emx2-CreERt2 mouse line production

A DNA sequence containing P2A and CreERt2 flanked by 800bp of homologous DNA was inserted into the third exon of *Emx2* using the Easi-Crispr strategy (Quadros, Miura et al. 2017) developed in conjunction with the University of Utah Mutation Generation and Detection Core facility. This gene targeting strategy replaces the endogenous stop codon of *Emx2* with P2A, and was directed by a guide RNA (gRNA) targeting the genomic sequence ‘CCTCAGACGATTAAAAGTCCAAA’. Double-stranded donor DNA containing the P2A-CreERt2 construct and flanking homologous arms was co-injected with gRNA and Cas9 protein by the University of Utah Transgenic and Gene Targeting Core facility. Accurate homologous recombination was confirmed by PCR amplification across the 5’ and 3’ homologous arms using primers binding to the *CreERt2* sequence (Fig.3, primer b (5’-CCGCCGCATAACCAGTGAAACAGC-3’) or primer c (5’-TGGCCCAGCTCCTCCTCATCCTCT-3’)) or the *Emx2* locus outside of either arm (Fig.3, primer a (5’-GGACGGGAGCAGAGGAAAGAGACC-3’) or primer d (5’-CCATCCCAGTCCTGCTCCCTCATT-3’)). Using this screening approach, 26% of progeny were identified that demonstrated *CreERt2* insertion at the *Emx2* allele. Three male founders (#s 2,15 and 45) with accurate homologous recombination within the 5’ and 3’ arms were further tested for Cre-recombinase activity resembling that of *Emx2*-Cre following tamoxifen induction at E15.5. A single male (#2) was selected and backcrossed with B6129S2/J F1 hybrid female mice to establish the *Emx2-*CreERt2 transgenic line.

### Tamoxifen induction

For CreER-mediated lineage tracing experiments, CreERt2 was induced by a single dose of 3mg Tamoxifen (Sigma Aldrich, T5648) and 3mg Progesterone (Sigma Aldrich, P3972) prepared in corn oil (Sigma Aldrich, C8267), and delivered via oral gavage to timed pregnant dams. Progesterone was included to reduce the likelihood of litter miscarriage due to tamoxifen toxicity. Pups were collected at P0 or gestational day 19.5 for dams that had not delivered litters before this stage.

### Immunofluorescent labeling and quantification

For wholemount immunofluorescent labeling heads were bisected and brains removed prior to immersion fixation of the intact inner ear and skull in 4% paraformaldehyde (PFA) solution (Electron Microscopy Sciences) prepared in 67mM Sorensons phosphate buffer (pH 7.4). Tissue was dissected to isolate the cochlea and vestibular organs and these were permeabilized for 30 minutes in blocking solution containing PBS with 1% bovine serum albumin (BSA) (Jackson ImmunoResearch 000-001-162) and 5% donkey serum (Jackson ImmunoResearch 017-000-121), supplemented with 0.5% Triton X-100 (Sigma Aldrich T9284). Primary antibodies were diluted in blocking solution supplemented with 0.1% Tween-20 (Sigma Aldrich 274348) and incubated with tissues overnight at 4°C. Tissue was rinsed thoroughly with PBS containing 0.05% Tween-20, and incubated overnight with species-specific AlexaFluor-conjugated secondary antibodies (Jackson ImmunoResearch 712-165-153, 705-545-147, 711-545-152, 711-605-152, 705-605-147, 711-165-152, 715-165-150; all used at a 1:1500 dilution).

Tissue was rinsed thoroughly with PBS containing 0.05% Tween-20 and mounted on slides using Prolong Gold (Thermo Fisher Scientific, P36930). For immunofluorescent labeling of cryosections, tissue was fixed with 4% PFA/Sorensons67 overnight at 4°C (P0 heads), or two hours on ice (embryos), then passed through a sucrose gradient and frozen in Neg-50 (Epredia). Sections were collected using a Leica CM3050 cryostat onto SuperFrost Plus slides (Fisher). Sections were labeled using a modified labeling protocol that excluded Tween-20 from the blocking and wash solutions. Fluorescent images were acquired by structured illumination microscopy using the Carl Zeiss Axio Imager M.2 with ApoTome.2 attachment and Axiocam 506 mono camera. Images were processed using Carl Zeiss Zen software, and figures prepared using Adobe Illustrator. FIJI(ImageJ) was used for quantification of tdTomato labeled cells in in the sensory and non-sensory regions of the vestibular organs. Commercial antibodies used in this study were: mouse anti-BetaIISpectrin (BD Biosciences 612562, 1:1000), rabbit anti-dsRed (Clontech/Takara 632496, 1:5000), rabbit anti-MyosinVIIA (Proteus Biosciences 25-6970, 1:800), goat anti-Oncomodulin (Santa Cruz Biotechnology SC-7466, 1:250), goat anti-Prox1 (R&D Systems AF2727, 1:300), goat anti-Sox2(Santa Cruz 17320, 1:200 for wholemount or 1:100 for cryosection), rat anti-tdTomato (Kerafast EST203, 1:500 for wholemount or 1:250 for cryosection), and Phalloidin Alexa Fluor 488 (Invitrogen, A12379, 1:1500).

## Notes

### Competing Interest Statement

The authors have declared no competing interest.

## References

Bermingham, N. A., B. A. Hassan, S. D. Price, M. A. Vollrath, N. Ben-Arie, R. A. Eatock, H. J. Bellen, A. Lysakowski and H. Y. Zoghbi (1999). “Math1: an essential gene for the generation of inner ear hair cells.” Science 284(5421): 1837–1841.

Bishop, K. M., G. Goudreau and D. D. O’Leary (2000). “Regulation of area identity in the mammalian neocortex by Emx2 and Pax6.” Science 288(5464): 344–349.

Bok, J., W. Chang and D. K. Wu (2007). “Patterning and morphogenesis of the vertebrate inner ear.” Int J Dev Biol 51(6-7): 521–533.

Bok, J., D. K. Dolson, P. Hill, U. Ruther, D. J. Epstein and D. K. Wu (2007). “Opposing gradients of Gli repressor and activators mediate Shh signaling along the dorsoventral axis of the inner ear.” Development 134(9): 1713–1722.

Deans, M. R. (2013). “A balance of form and function: planar polarity and development of the vestibular maculae.” Semin Cell Dev Biol 24(5): 490–498.

Deans, M. R. (2021). “Conserved and Divergent Principles of Planar Polarity Revealed by Hair Cell Development and Function.” Front Neurosci 15: 742391.

Deans, M. R., D. Antic, K. Suyama, M. P. Scott, J. D. Axelrod and L. V. Goodrich (2007). “Asymmetric distribution of prickle-like 2 reveals an early underlying polarization of vestibular sensory epithelia in the inner ear.” J Neurosci 27(12): 3139–3147.

Fritzsch, B., K. W. Beisel, K. Jones, I. Farinas, A. Maklad, J. Lee and L. F. Reichardt (2002). “Development and evolution of inner ear sensory epithelia and their innervation.” J Neurobiol 53(2): 143–156.

Ghimire, S. R., E. M. Ratzan and M. R. Deans (2018). “A non-autonomous function of the core PCP protein VANGL2 directs peripheral axon turning in the developing cochlea.” Development 145(12).

Goodrich, L. V. (2016). Early Development of the Spiral Ganglion. The Primary Auditory Neurons of the Mammalian Cochlea. A. F. Dabdoub, B.; Popper, A.N.; Fray, R.R.. New York, Springer Science+Business Media. 52: 11–48.

Gu, R., R. M. Brown, 2nd, C. W. Hsu, T. Cai, A. L. Crowder, V. G. Piazza, T. J. Vadakkan, M. E. Dickinson and A. K. Groves (2016). “Lineage tracing of Sox2-expressing progenitor cells in the mouse inner ear reveals a broad contribution to non-sensory tissues and insights into the origin of the organ of Corti.” Dev Biol 414(1): 72–84.

Hamasaki, T., A. Leingartner, T. Ringstedt and D. D. O’Leary (2004). “EMX2 regulates sizes and positioning of the primary sensory and motor areas in neocortex by direct specification of cortical progenitors.” Neuron 43(3): 359–372.

Holley, M., C. Rhodes, A. Kneebone, M. K. Herde, M. Fleming and K. P. Steel (2010). “Emx2 and early hair cell development in the mouse inner ear.” Dev Biol 340(2): 547–556.

Ji, Y. R., Y. Tona, T. Wafa, M. E. Christman, E. D. Tourney, T. Jiang, S. Ohta, H. Cheng, T. Fitzgerald, B. Fritzsch, S. M. Jones, K. E. Cullen and D. K. Wu (2022). “Function of bidirectional sensitivity in the otolith organs established by transcription factor Emx2.” Nat Commun 13(1): 6330.

Ji, Y. R., S. Warrier, T. Jiang, D. K. Wu and K. S. Kindt (2018). “Directional selectivity of afferent neurons in zebrafish neuromasts is regulated by Emx2 in presynaptic hair cells.” Elife 7.

Jia, S., E. M. Ratzan, E. J. Goodrich, R. Abrar, L. Heiland, B. Tarchini and M. R. Deans (2023). “The dark kinase STK32A regulates hair cell planar polarity opposite of EMX2 in the developing mouse inner ear.” Elife 12.

Jiang, T., K. Kindt and D. K. Wu (2017). “Transcription factor Emx2 controls stereociliary bundle orientation of sensory hair cells.” Elife 6.

Keithley, E. M., L. Erkman, T. Bennett, L. Lou and A. F. Ryan (1999). “Effects of a hair cell transcription factor, Brn-3.1, gene deletion on homozygous and heterozygous mouse cochleas in adulthood and aging.” Hear Res 134(1-2): 71–76.

Kimura, J., Y. Suda, D. Kurokawa, Z. M. Hossain, M. Nakamura, M. Takahashi, A. Hara and S. Aizawa (2005). “Emx2 and Pax6 function in cooperation with Otx2 and Otx1 to develop caudal forebrain primordium that includes future archipallium.” J Neurosci 25(21): 5097–5108.

Kindt, K. S., A. Akturk, A. Jarysta, M. Day, A. Beirl, M. Flonard and B. Tarchini (2021). “EMX2-GPR156-Galphai reverses hair cell orientation in mechanosensory epithelia.” Nat Commun 12(1): 2861.

Koo, S. K., J. K. Hill, C. H. Hwang, Z. S. Lin, K. J. Millen and D. K. Wu (2009). “Lmx1a maintains proper neurogenic, sensory, and non-sensory domains in the mammalian inner ear.” Dev Biol 333(1): 14–25.

Kozak, E. L., S. Palit, J. R. Miranda-Rodriguez, A. Janjic, A. Bottcher, H. Lickert, W. Enard, F. J. Theis and H. Lopez-Schier (2020). “Epithelial Planar Bipolarity Emerges from Notch-Mediated Asymmetric Inhibition of Emx2.” Curr Biol 30(6): 1142–1151 e1146.

Lozano-Ortega, M., G. Valera, Y. Xiao, A. Faucherre and H. Lopez-Schier (2018). “Hair cell identity establishes labeled lines of directional mechanosensation.” PLoS Biol 16(7): e2004404.

Madisen, L., T. A. Zwingman, S. M. Sunkin, S. W. Oh, H. A. Zariwala, H. Gu, L. L. Ng, R. D. Palmiter, M. J. Hawrylycz, A. R. Jones, E. S. Lein and H. Zeng (2010). “A robust and high-throughput Cre reporting and characterization system for the whole mouse brain.” Nat Neurosci 13(1): 133–140.

Mallamaci, A., L. Muzio, C. H. Chan, J. Parnavelas and E. Boncinelli (2000). “Area identity shifts in the early cerebral cortex of Emx2-/- mutant mice.” Nat Neurosci 3(7): 679–686.

Mann, Z. F., H. Galvez, D. Pedreno, Z. Chen, E. Chrysostomou, M. Zak, M. Kang, E. Canden and N. Daudet (2017). “Shaping of inner ear sensory organs through antagonistic interactions between Notch signalling and Lmx1a.” Elife 6.

Morsli, H., D. Choo, A. Ryan, R. Johnson and D. K. Wu (1998). “Development of the mouse inner ear and origin of its sensory organs.” J Neurosci 18(9): 3327–3335.

Nichols, D. H., S. Pauley, I. Jahan, K. W. Beisel, K. J. Millen and B. Fritzsch (2008). “Lmx1a is required for segregation of sensory epithelia and normal ear histogenesis and morphogenesis.” Cell Tissue Res 334(3): 339–358.

Ono, K., T. Kita, S. Sato, P. O’Neill, S. S. Mak, M. Paschaki, M. Ito, N. Gotoh, K. Kawakami, Y. Sasai and R. K. Ladher (2014). “FGFR1-Frs2/3 signalling maintains sensory progenitors during inner ear hair cell formation.” PLoS Genet 10(1): e1004118.

Quadros, R. M., H. Miura, D. W. Harms, H. Akatsuka, T. Sato, T. Aida, R. Redder, G. P. Richardson, Y. Inagaki, D. Sakai, S. M. Buckley, P. Seshacharyulu, S. K. Batra, M. A. Behlke, S. A. Zeiner, A. M. Jacobi, Y. Izu, W. B. Thoreson, L. D. Urness, S. L. Mansour, M. Ohtsuka and C. B. Gurumurthy (2017). “Easi-CRISPR: a robust method for one-step generation of mice carrying conditional and insertion alleles using long ssDNA donors and CRISPR ribonucleoproteins.” Genome Biol 18(1): 92.

Rhodes, C. R., N. Parkinson, H. Tsai, D. Brooker, S. Mansell, N. Spurr, A. J. Hunter, K. P. Steel and S. D. Brown (2003). “The homeobox gene Emx2 underlies middle ear and inner ear defects in the deaf mouse mutant pardon.” J Neurocytol 32(9): 1143–1154.

Steffes, G., B. Lorente-Canovas, S. Pearson, R. H. Brooker, S. Spiden, A. E. Kiernan, J. L. Guenet and K. P. Steel (2012). “Mutanlallemand (mtl) and Belly Spot and Deafness (bsd) are two new mutations of Lmx1a causing severe cochlear and vestibular defects.” PLoS One 7(11): e51065.

Stoller, M. L., O. Roman, Jr. and M. R. Deans (2018). “Domineering non-autonomy in Vangl1;Vangl2 double mutants demonstrates intercellular PCP signaling in the vertebrate inner ear.” Dev Biol 437(1): 17–26.

Wallis, D., M. Hamblen, Y. Zhou, K. J. Venken, A. Schumacher, H. L. Grimes, H. Y. Zoghbi, S. H. Orkin and H. J. Bellen (2003). “The zinc finger transcription factor Gfi1, implicated in lymphomagenesis, is required for inner ear hair cell differentiation and survival.” Development 130(1): 221–232.

Whitfield, T. T. and K. L. Hammond (2007). “Axial patterning in the developing vertebrate inner ear.” Int J Dev Biol 51(6-7): 507–520.

Yang, X., X. Qian, R. Ma, X. Wang, J. Yang, W. Luo, P. Chen, F. Chi and D. Ren (2017). “Establishment of planar cell polarity is coupled to regional cell cycle exit and cell differentiation in the mouse utricle.” Sci Rep 7: 43021.

Zak, M. and N. Daudet (2021). “A gradient of Wnt activity positions the neurosensory domains of the inner ear.” Elife 10.

